# Topoisomerase I promotes formation of a novel DNA intermediate in the ribosomal DNA

**DOI:** 10.1101/2022.05.24.493248

**Authors:** Temistocles Molinar, Daniel Sultanov, Hannah Klein, Andreas Hochwagen

## Abstract

The ribosomal DNA (rDNA) is a conserved but highly unstable multi-gene locus that codes for the RNA components of the ribosome (rRNA). Despite being essential for protein translation and survival, rRNA gene-copy numbers fluctuate frequently. Here, we describe a novel rDNA intermediate that may be involved in these copy-number changes in *Saccharomyces cerevisiae*. Two-dimensional gel analyses revealed a structure that includes a large single-stranded tail and arises from the promoter of the 35S rRNA gene. Formation of this intermediate is unaffected by reduced 35S transcription but relies on topoisomerase I (Top1). Unexpectedly, intermediate formation is also independent of S phase or the rDNA-instability factor Fob1, even though Fob1 is mediates the major peak of Top1 cleavage complexes in the rDNA. Indeed, we find that the known rDNA instability phenotypes of *top1* mutants, including increased formation of extrachromosomal rDNA circles, elevated genetic marker loss, and instability of critically short rDNA arrays, are also largely independent of Fob1. We therefore speculate that failure to form this intermediate leads to rDNA instability in *top1* mutants.

## INTRODUCTION

The RNA components of the ribosome (rRNAs) are essential for the catalytic activity and structural integrity of the ribosome and as such are required for viability in all forms of life. To support the high demand for ribosomes, rRNA genes are typically encoded in one or several multigene loci, predisposing the ribosomal DNA (rDNA) to instability caused by non-allelic homologous recombination. Consequently, rDNA copy numbers can fluctuate widely even from one generation to the next (Stults et al., 2008), and spontaneous loss events that eliminate many copies are common (Michel et al., 2005, Ganley and Kobayashi, 2011).

Several mechanisms help cells to increase rDNA copy number and regrow shortened rDNA arrays. In yeast, the replication fork-barrier protein Fob1 simulates the formation of DNA breaks whose repair by unequal sister chromatid exchange changes rDNA copy number (Kobayashi, 2011). Genetically encoded replication fork barriers are observed in the rDNA of many eukaryotes, although available evidence suggests that Fob1 activates rDNA recombination independently of its fork-barrier activity (Gadaleta and Noguchi, 2017, Ward et al., 2000, Krawczyk et al., 2014). Fob1 also promotes the replicative production of extra-chromosomal rDNA circles (ERCs), which can insert into the rDNA (Mansisidor et al., 2018). Additionally, in the absence of the histone chaperone Asf1, rDNA arrays grow independently of Fob1, using a mechanism that may involve a form of break-induced replication (Houseley and Tollervey, 2011).

In yeast, the topoisomerase Top1 has an important but poorly understood role in controlling rDNA copy number and stability. Top1 is a type I topoisomerase that reversibly cuts one strand of the DNA backbone to allow DNA swiveling and relaxation of DNA supercoils. Top1 is enriched in the nucleolus, the subnuclear structure organized by the rDNA and shows two peaks of enrichment within rDNA sequences: a major Fob1-dependent peak at the replication fork barrier and a smaller Fob1-independent peak in the promoter of the 35S rRNA gene (Krawczyk et al., 2014, Di Felice et al., 2019). Cleavage complexes with Top1 covalently attached to the DNA substrate indicate that Top1 catalyzes strand breakage at both locations. Top1-dependent nicking at the replication fork barrier has been proposed to mediate the creation of a single-ended double-strand break (DSB) upon DNA replication that may stimulate rDNA recombination (Krawczyk et al., 2014). In absence of Top1, the rDNA array becomes unstable and ERC levels are increased (Krawczyk et al., 2014, Kim and Wang, 1989, Christman et al., 1988), possibly because of topological stress from unresolved supercoils. However, mutants with defective topoisomerase II (Top2) show a distinct phenotype of rDNA instability (Christman et al., 1993), implying that Top1 has roles in the rDNA that cannot be compensated by Top2.

Here, we report that Top1 promotes the formation a novel DNA intermediate that originates from the promoter of the 35S rRNA gene in *Saccharomyces cerevisiae*. The intermediate contains a long single-stranded tail and forms independently of S phase or Fob1. Failure to form this intermediate is associated with substantial Fob1-independent rDNA instability in *top1* mutants.

## RESULTS

### An rDNA intermediate with a long single-stranded tail

To investigate possible repair intermediates associated with fluctuating rDNA copy numbers, we analyzed the rDNA of cycling wild-type cells using two-dimensional gel electrophoresis and Southern blotting. This technique separates DNA fragments based on both size and shape and has been used extensively to study replication intermediates in the rDNA (Brewer and Fangman, 1987, Brewer and Fangman, 1988). When we analyzed a StuI fragment that spanned the intergenic spacer regions (*IGS1* and *IGS2*) as well as part of the 35S gene, we not only detected the expected signals of replication forks and the fork barrier but also a novel fast-migrating arc (**Figure 1a**). The arc intersected with the sheared DNA below the linear restriction fragment and extended upward, indicating that the arc represents a series of DNA species with progressively higher molecular weights.

**Figure 1.**
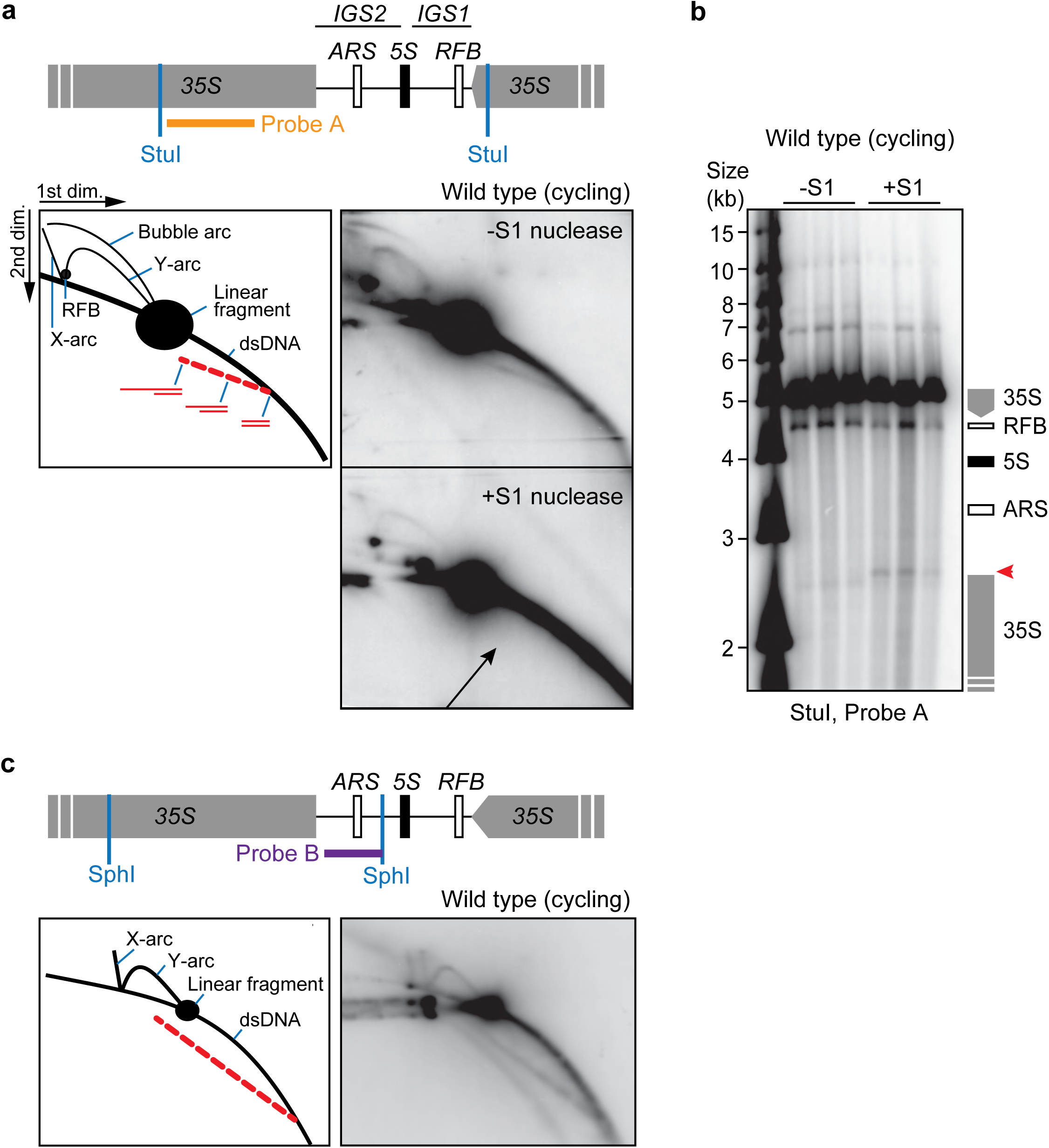
An ssDNA-containing rDNA intermediate originating from the 35S promoter. **a**. Two-dimensional gel electrophoresis and Southern analysis of a StuI fragment encompassing the intergenic regions and part of the 35S rRNA gene isolated from logarithmically cycling wild-type cells. Schematic on top shows locations of restriction sites and probe relative to rDNA landmarks, including the origin of replication (ARS), the replication fork barrier (RFB) and the 5S and 35S rRNA genes. Schematic in left box indicates the positions of replication-dependent structures (Y-arc, bubble arc, X-arc and RFB) above the linear restriction fragment as well as a novel arc (red dashed line), which likely represents an intermediate with a single-stranded tail of increasing length. Boxes on the right show the same samples but bottom sample was incubated with S1 nuclease prior to electrophoresis, leading to a disappearance of the novel arc (arrow). **b**. One-dimensional gel electrophoresis of the same StuI fragment shows that S1 treatment leads to the appearance of a new band mapping near the 35S promoter (red arrowhead). Schematic on the right shows the relative positions of rDNA landmarks. Each lane is an independent replicate DNA sample. **c**. Two-dimensional gel electrophoresis and Southern analysis of an SphI fragment encompassing part of the intergenic regions and and 35S rRNA gene. RFB signal and bubble arc are not seen because the RFB is located in the adjacent fragment and the origin is toward one end of the fragment, but the novel arc is clearly detectable (dashed red line in schematic on the left).

The fast migration of the arc, ahead of linear double-stranded DNA, suggested that it contained single-stranded DNA (ssDNA). Consistent with this interpretation, the arc became undetectable when samples were treated with S1 nuclease prior to electrophoresis (**Figure 1a**). Given that the arc also intersected with signals from linear double-stranded DNA, the arc likely represents a double-stranded fragment of uniform size with a variably sized single-stranded tail that can reach considerable lengths. Most likely, the increasing molecular weight of the arc is the result of DNA synthesis, although the arc could theoretically also arise from distributed DNA breakage followed by resection up to a defined point. The arc is unlikely to contain RNA because samples were incubated overnight with RNase A as part of the DNA extraction procedure (see Methods).

To more precisely map the point of origin of the arc, we analyzed S1-nuclease-treated samples by standard gel electrophoresis. This analysis revealed a distinct new band in S1 nuclease-treated samples (**Figure 1b**), supporting the idea that the different ssDNA-containing species that form the arc originate from a single point. Based on the size of the new band, this point is at or near the promoter region of the *35S* pre-rRNA gene. To further support this interpretation, we performed 2D gel analyses using several different restriction enzymes and probes. We only observed the arc in restriction fragments that centered on the 35S promoter (**Figure 1c** and data not shown). These data indicate that the 35S promoter region gives rise to a DNA species with a long single-stranded tail.

### The intermediate is independent of S phase

To test whether the tailed intermediate is linked to S-phase-dependent DNA synthesis, we enriched cells in G1 using alpha-factor. Accumulation of cells in G1 strongly reduced the signal intensity of replication-dependent bubble- and Y-arcs but did not decrease the intensity of the tailed intermediate, indicating that the intermediate forms independently of canonical replication structures (**Figure 2a**). To further support this conclusion, we followed rDNA replication in a synchronized culture. Cells were arrested in telophase using a temperature-sensitive *cdc15-2* mutation and released into G1 by shifting cells to the permissive temperature (Futcher, 1999). DNA replication intermediates became apparent 45 minutes after the release, and were maximal at 60 minutes before declining again (**Figure 2b**), in line with the expected S-phase kinetics (Futcher, 1999). By contrast, the tailed intermediate was detectable at all time points, albeit with somewhat varying signal intensities. We conclude that formation of the tailed intermediate does not depend on S phase.

**Figure 2.**
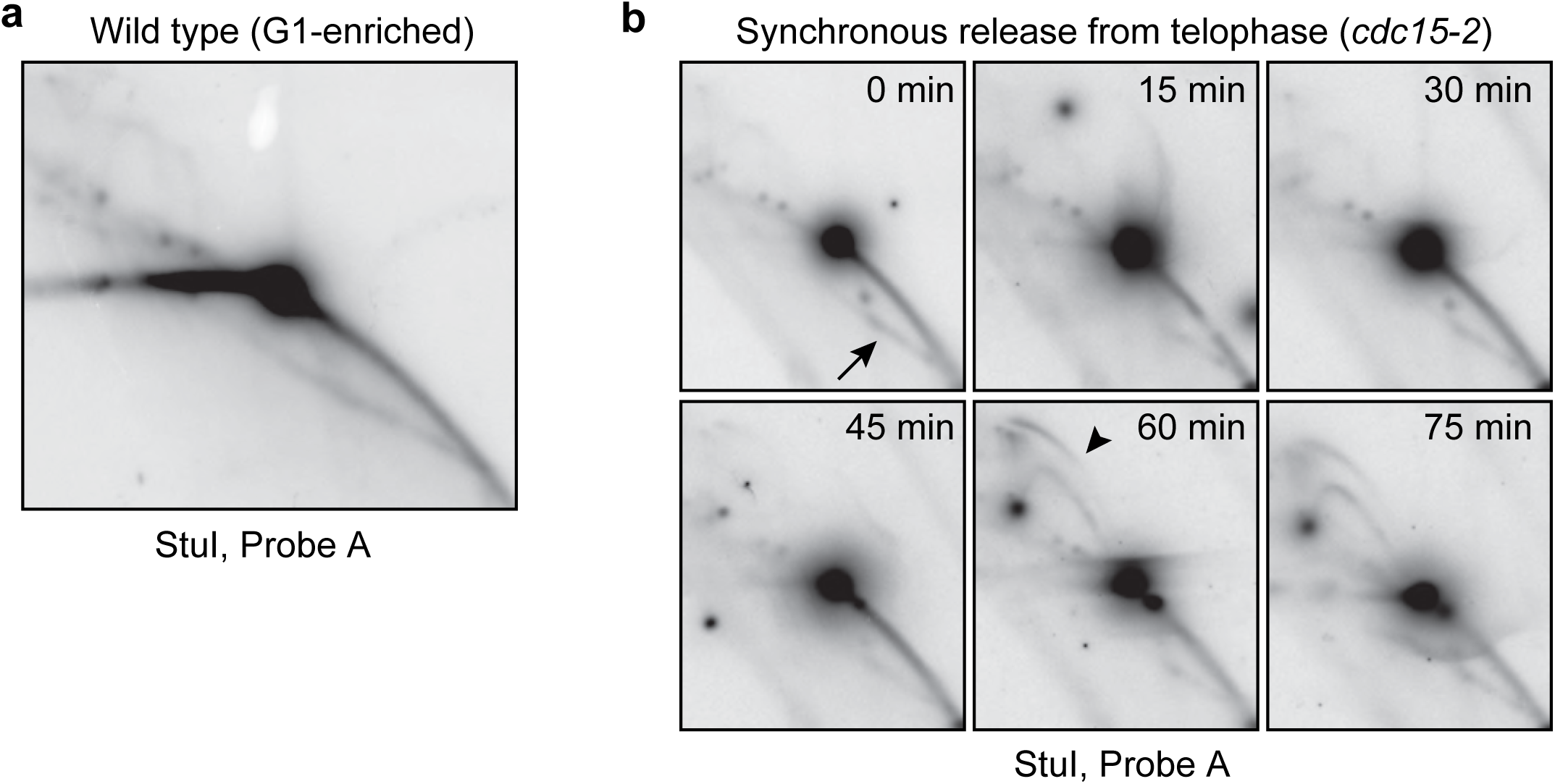
The tailed intermediate forms independently of S phase. **a**. Two-dimensional gel electrophoresis and Southern analysis of a StuI fragment isolated from cells that were accumulated in G1 using alpha-factor pheromone. Replication intermediates are nearly undetectable but the tailed intermediate is prominently visible. **b**. Time course of *cdc15-2* cells progressing synchronously from telophase upon shift to the permissive temperature. The tailed intermediate (arrow) is present throughout the time course, whereas replication intermediates (arrowhead) are undetectable for the first 30 minutes and are maximal 60 minutes after release from telophase.

### Formation of the tailed intermediate requires topoisomerase I

We sought to identify factors required to produce the tailed intermediate. The origin of the ssDNA at or near the 35S promoter suggested a possible role for transcriptional activity. Therefore, we analyzed strains lacking the non-essential RNA polymerase I (RNAPI) subunit Rpa49, the RNAPI transcription factor Uaf30, and the high-mobility group protein Hmo1, all of which have reduced 35S transcription (Gadal et al., 2002, Beckouet et al., 2008, Siddiqi et al., 2001). However, in all three mutants, the tailed intermediate was still clearly detectable (**Figure 3a-b**). These data indicate that formation of the tailed intermediate does not require strong transcription of the 35S transcription units.

**Figure 3.**
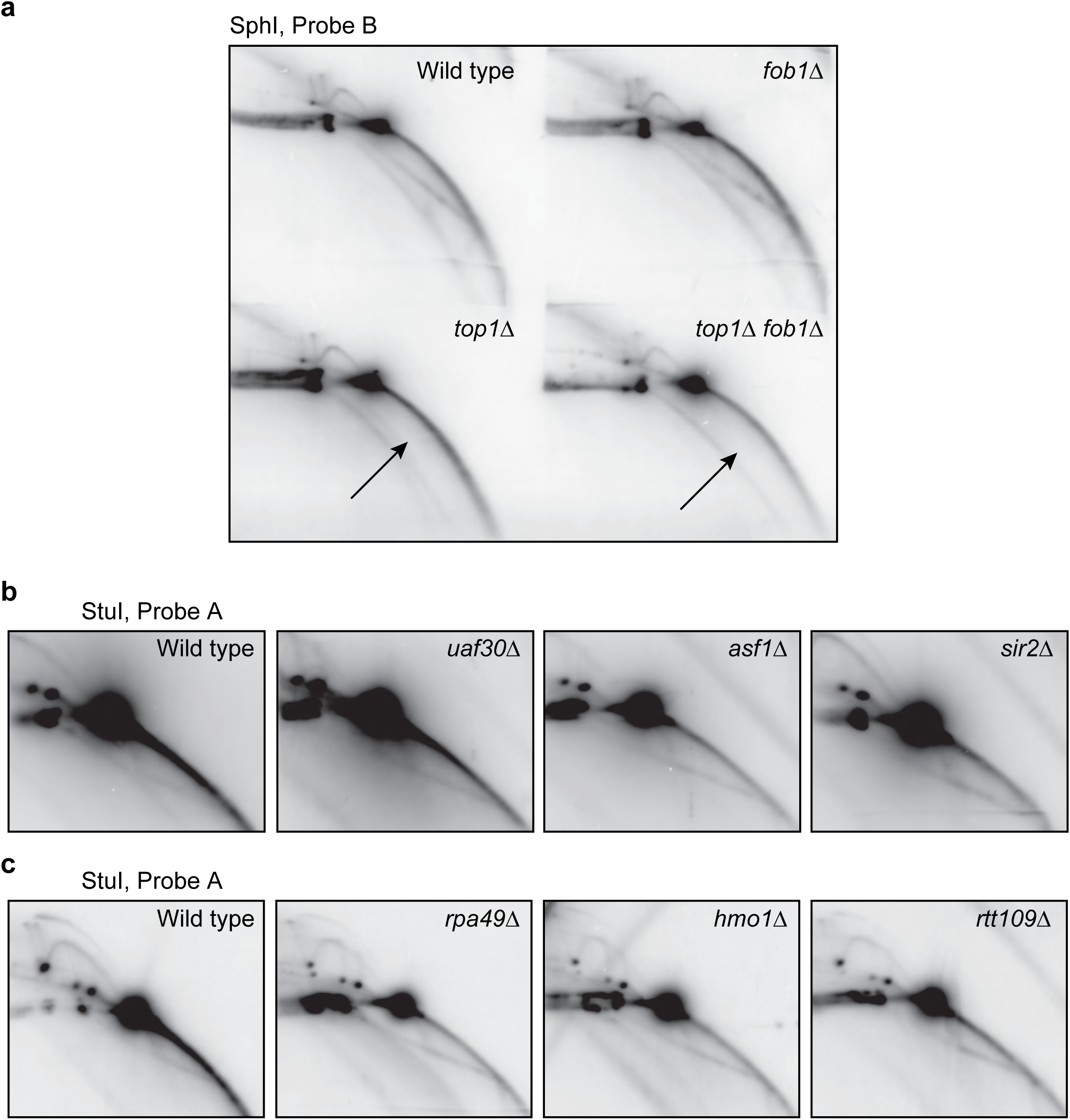
The tailed intermediate requires Top1 but is independent of Fob1 or full 35S transcription. **a**. Two-dimensional gel electrophoresis and Southern analysis of an SphI rDNA fragment from wild type, *fob1Δ, top1Δ* or *fob1Δ top1Δ* logarithmic cultures. Arrows indicate the absence of the tailed intermediate in *top1Δ* and *fob1Δ top1Δ* samples. **b**. and **c**. Two-dimensional gel electrophoresis and Southern analysis of a StuI rDNA fragment in mutants lacking regulators of rDNA transcription and stability.

The promoter of the *35S* pre-rRNA gene is also a site of transcription-independent cleavage activity by topoisomerase I (Vogelauer and Camilloni, 1999, Krawczyk et al., 2014, Di Felice et al., 2019). Indeed, the tailed intermediate was undetectable when we analyzed *top1* mutants using different restriction fragments and probes (**Figure 3c, Supplemental Figure 1**). We conclude that Top1 is essential for the formation of the tailed intermediate.

Previous analyses defined a recombination-stimulating activity of Fob1 that requires both the fork barrier and the 35S promoter, and showed that Fob1 induces rDNA nicking independently of S phase (Voelkel-Meiman et al., 1987, Lin and Keil, 1991, Kobayashi et al., 1998, Burkhalter and Sogo, 2004). Therefore, we also analyzed *fob1* mutants. However, formation of the tailed intermediate appeared unaffected in *fob1* mutants and only became undetectable when Top1 was also removed (**Figure 3c**). These data indicate that formation of the intermediate occurs independently of the fork barrier activity in the rDNA, and thus does not require one of the major known drivers of rDNA instability.

The intermediate was also formed normally in several mutants with pronounced Fob1-dependent or Fob1-independent rDNA instability, including mutants lacking the histone deacetylase Sir2, the histone H3K56 acetyltransferase Rtt109, and the histone chaperone Asf1 (Kaeberlein et al., 1999, Ide et al., 2013, Houseley and Tollervey, 2011) (**Figure 3b-c**). These data imply that the intermediate is likely the product of a specific form of rDNA metabolism, rather than a general consequence of rDNA instability.

### rDNA instability of *top1* mutants is independent of Fob1

*top1* mutants exhibit profound rDNA instability (Krawczyk et al., 2014, Kim and Wang, 1989, Christman et al., 1988). Although this instability was proposed to be linked to the Fob1-dependent recruitment of Top1 to the replication fork barrier, this connection remained untested (Krawczyk et al., 2014). Our results alternatively suggested that the rDNA instability of *top1* mutants might be linked to the inability to form the tailed intermediate. If so, then the instability phenotypes of *top1* mutants, like the tailed intermediate, should be independent of Fob1.

We used several approaches to test this prediction. Pulsed-field gel electrophoresis analysis of chromosomal rDNA fragments from *top1* mutants revealed the expected smearing, as well as increased amounts of rDNA-containing sequences in in the wells, indicating the presence of branched DNA structures or rDNA circles that failed to enter the gel (Houseley et al., 2007, Krawczyk et al., 2014, Christman et al., 1988) (**Figure 4a**). However, both, smearing and retention in the wells were largely independent of Fob1. Similarly, the high ERC levels seen in *top1* mutants (Krawczyk et al., 2014, Kim and Wang, 1989) were only mildly reduced by deleting *FOB1* (**Figure 4b**). These data show that the rDNA instability of *top1* mutants is largely independent of Fob1 activity. To confirm this observation, we monitored the stability of a *URA3* marker inserted into the rDNA. Loss of Top1 led to the expected increase in marker loss from the rDNA (Christman et al., 1988), but these loss events were independent of Fob1 (**Figure 4c**). These data are consistent with the model that the Fob1-independent function of Top1 in forming the tailed rDNA intermediates may be important for rDNA stability.

**Figure 4.**
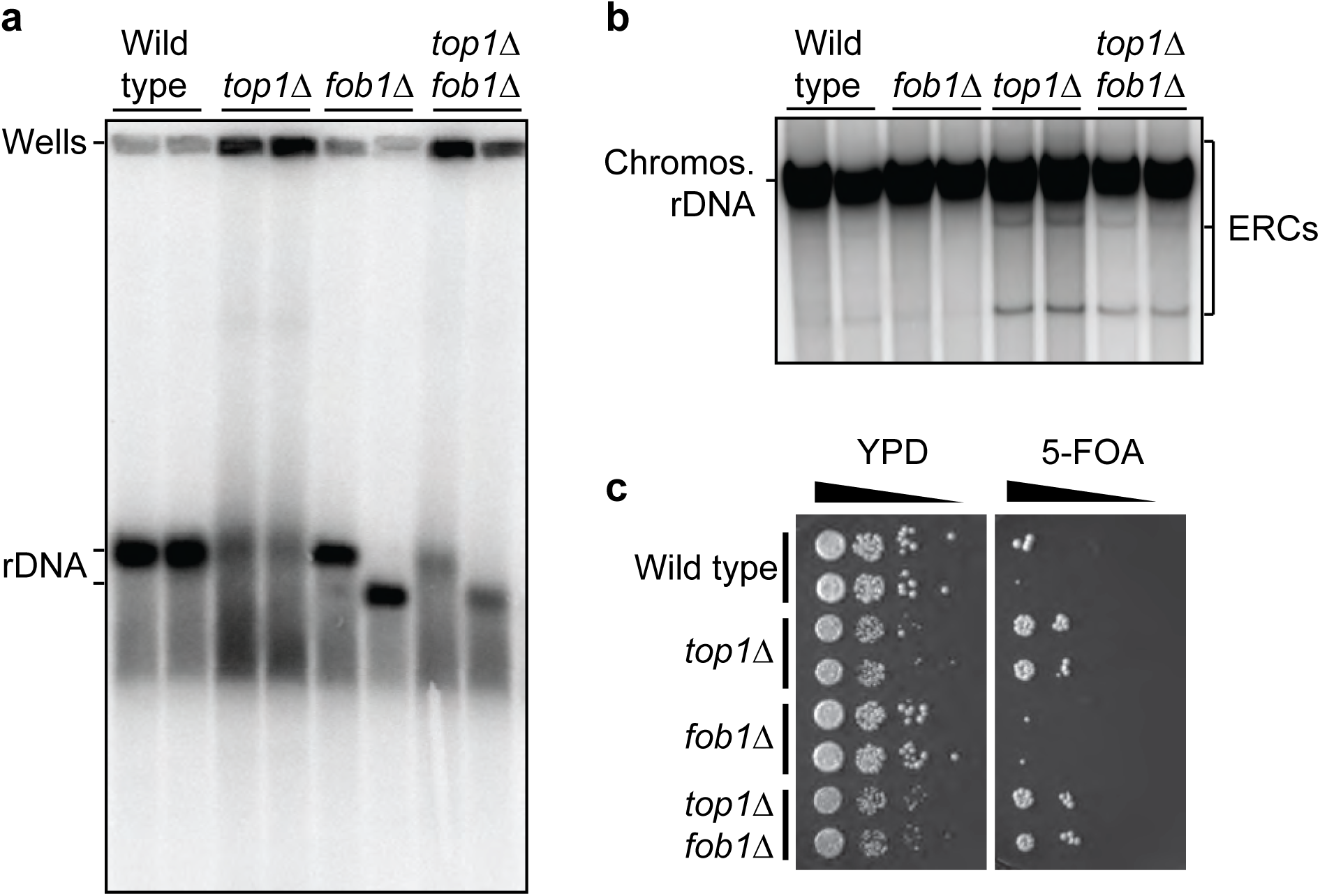
The rDNA instability of *top1* mutants is independent of Fob1. **a**. Pulsed-field gel electrophoresis and Southern analysis of entire rDNA arrays from wild type, *top1Δ, fob1Δ* or *top1Δ fob1Δ* logarithmic cultures. rDNA arrays were released from chromosome XII by digesting with XhoI, which cuts outside the rDNA but leaves the rDNA intact. Lanes show biological replicates. Wild type and *top1Δ* strains represent two sibling pairs; *fob1Δ* or *top1Δ fob1Δ* strains also represent two sibling pairs from single tetrads. The crosses used genetically tagged arrays in *trans* to ensure the same array was analyzed. **b**. Standard gel electrophoresis and Southern analysis of undigested genomic samples from wild type, *fob1Δ, top1Δ* or *fob1Δ top1Δ* logarithmic cultures. Circular species (ERCs) migrate slower and faster than the sheared chromosomal rDNA, which migrates as a single band in this analysis. Lanes are biological replicates. Probe B was used for both (a) and (b). **c**. Wild type, *top1Δ, fob1Δ* or *top1Δ fob1Δ* logarithmic cultures carrying a *URA3* marker in repeat 3 of the rDNA array were 10-fold serially diluted and spotted on rich medium (YPD) or medium or medium containing 5-fluoroorotic acid, which selects for cells that have lost the *URA3* marker. Rows are independent biological replicates.

### Critically short rDNA arrays undergo stepwise expansion

To further investigate the role of Top1 in preventing rDNA instability, we created critically short rDNA arrays and deleted *FOB1* to prevent Fob1-dependent re-expansion (Kobayashi, 2006, Kobayashi et al., 1998). Fine-scale copy-number analysis of such critically short rDNA arrays revealed an intriguing laddering pattern of instability that occurred despite the absence of Fob1, indicating that a Fob1-independent mechanism helps expand critically short rDNA arrays (**Figure 5a**). Laddering occurred in step sizes of ∼9 kb, matching the size of a single rDNA repeat, and exhibited a pronounced bias toward higher copy numbers that is likely driven by the slow growth phenotype of cells with critically short rDNA arrays (Takeuchi et al., 2003). In addition, larger jumps in copy number were apparent. Interestingly, induction of *FOB1* from an inducible promoter construct that recapitulates endogenous Fob1 levels (Mansisidor et al., 2018) did not alter the laddering over several population doublings, indicating that Fob1 does not promote rapid rDNA expansion in the short term.

**Figure 5.**
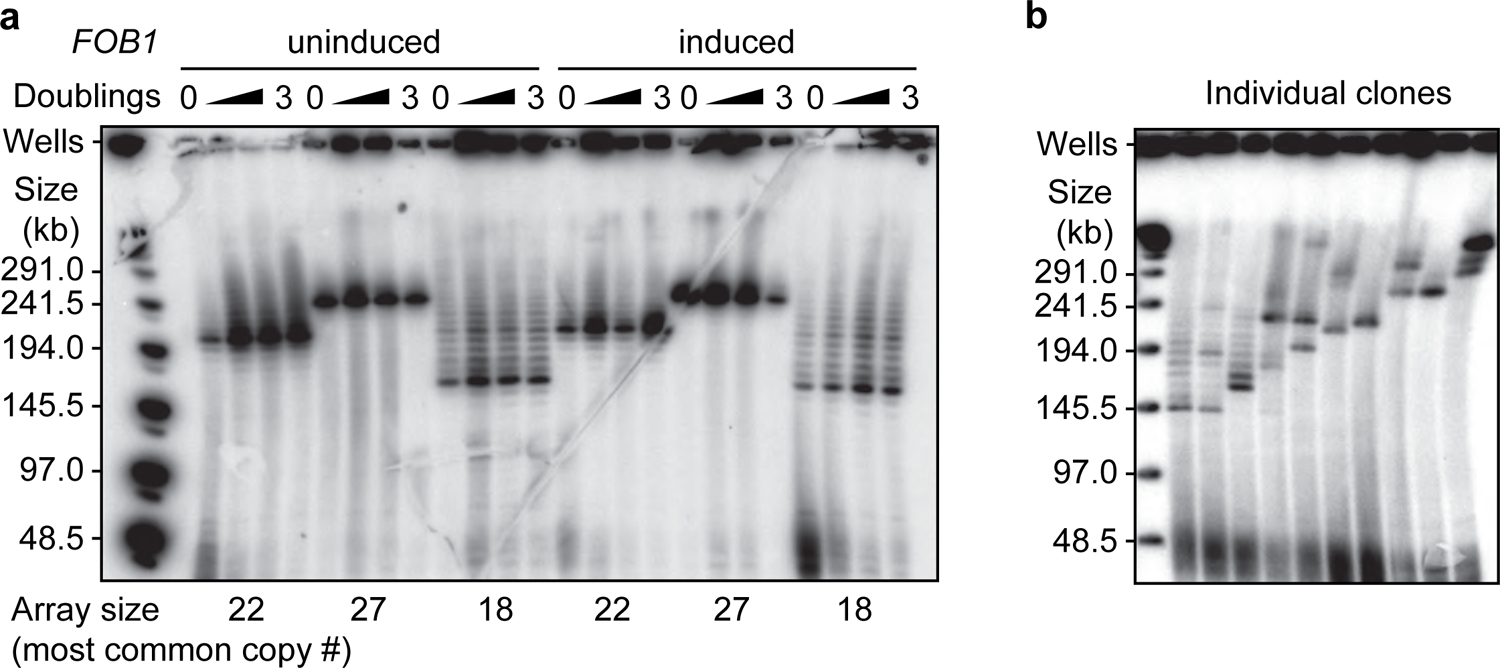
Critically short rDNA arrays undergo stepwise rDNA expansion independently of Fob1. **a**. Pulsed-field gel electrophoresis and Southern analysis of isolates with critically short rDNA arrays carrying a conditional *FOB1* construct that allows induction of wild-type Fob1 levels upon addition of β-estradiol (Mansisidor et al., 2018). Isolates were pre-grown in YPD before cultures were split and Fob1 induced in one half by β-estradiol addition. Culture density was monitored by spectrophotometer and samples were collected after each culture doubling for 3 doublings. Cells were embedded and in agarose plus and arrays released by XhoI digest before analysis. The array sizes noted below are based on the most prominent bands in the ladders. **b**. Cell clones were isolated from the 18-repeat culture in (a), expanded in YPD and analyzed by pulsed-field gel electrophoresis and Southern analysis.

To test whether the laddering reflects copy-number heterogeneity within the culture, we isolated and expanded individual clones of the culture and assessed their copy numbers. This analysis revealed a varied number of rDNA copies in the clones, with each clone again showing biased laddering toward higher copy number (**Figure 5b**). These data indicate that laddering is an ongoing process that creates copy-number heterogeneity in cultures with critically short rDNA arrays.

### Loss of Top1 causes instability of critically short rDNA arrays

To assess the effect of the *top1* mutation on critically short rDNA arrays, it was important to clearly distinguish *TOP1*-dependent effects from the inherent instability of critically short rDNA arrays. We achieved this by analyzing sister spores that inherited the same genetically marked rDNA array but differed in their inheritance of *TOP1* vs. *top1Δ* (**Figure 6a**; see Methods). In this way, the two siblings started with identical arrays and any differences in size were the consequence of events (*TOP1*-dependent or spontaneous) that occurred during the outgrowth of the sibling spores.

**Figure 6.**
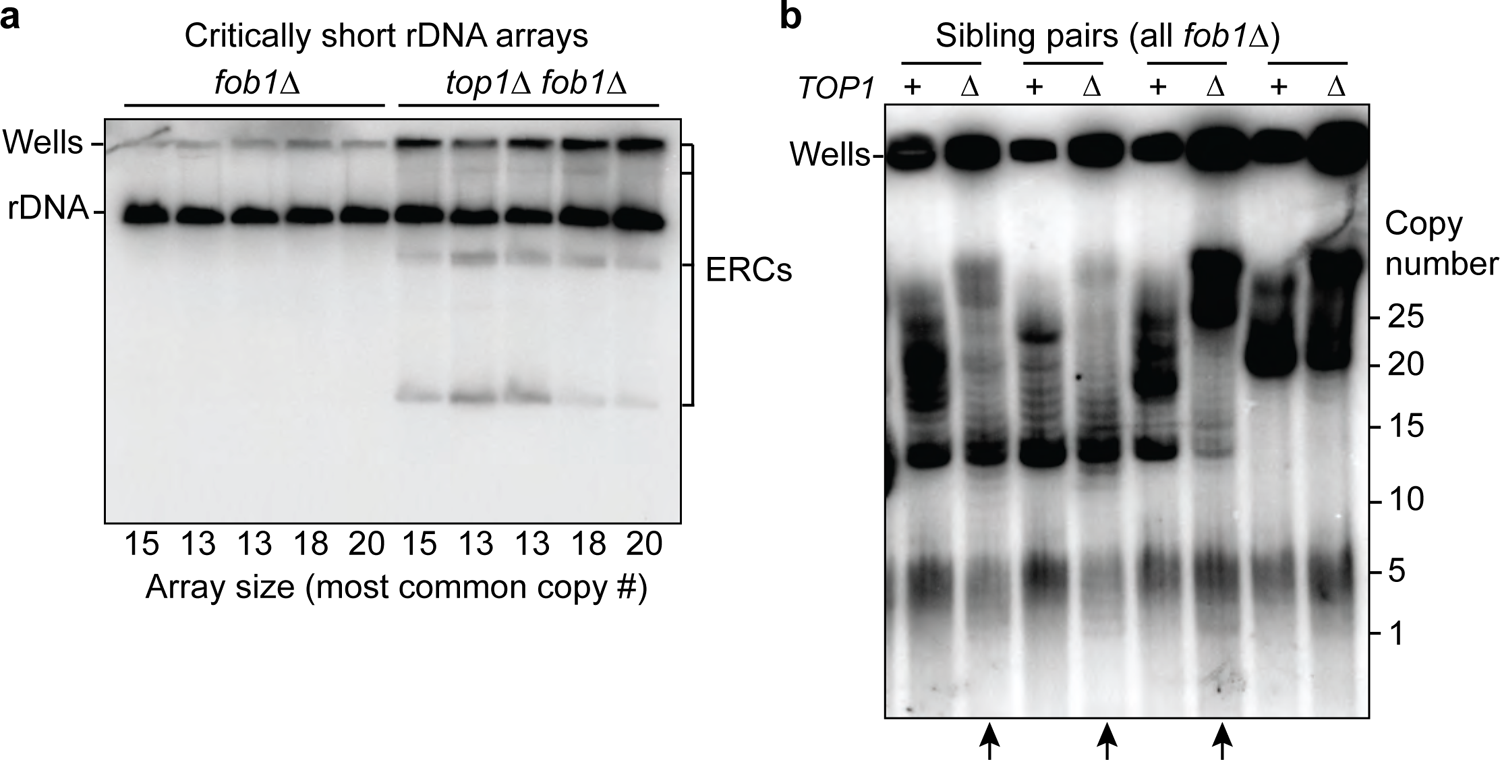
Increased instability of critically short rDNA arrays in the absence of topoisomerase I. **a**. Standard gel electrophoresis and Southern analysis of XhoI-digested genomic samples from wild-type and *top1Δ* sibling strains with genetically marked, short rDNA arrays grown in log phase (siblings are the offspring of a cross between AH11094 and AH11402 and are arranged in order for each genotype). All strains lack *FOB1*. The array sizes noted below are based on the most prominent bands in the ladders obtained from pulsed-field gel analyses. Circular species (ERCs) migrate slower and faster than the sheared chromosomal rDNA, which migrates as a single band. **b**. Pulsed-field gel electrophoresis and Southern analysis of critically short rDNA arrays from wild type/ *top1Δ* sibling pairs (all strains lack *FOB1*). rDNA arrays were released by XhoI digest prior to analysis. Pairs are arranged as neighbors. Arrows indicate lanes with increased laddering toward lower copy numbers.

The sibling analysis revealed several phenotypes caused by the loss of *TOP1*. First, ERC levels were strongly increased in *top1* siblings with critically short arrays despite the absence of Fob1 (**Figure 6a**). This increase is in line with the instability seen for larger arrays (**Figure 4b**). Second, *top1* siblings exhibited a faster increase in rDNA sizes than the corresponding wild-type siblings, as evident from the higher maximal rDNA copy numbers compared to *TOP1* siblings (**Figure 6b**). Thus, loss of *TOP1* causes less controlled rDNA expansion. Finally, *top1* siblings consistently also showed more laddering toward *lower* copy numbers. Indeed, in several siblings, laddering was apparent even down to single rDNA copies (**Figure 6b**; arrows), suggesting that the mode and/or frequency of recombination and rDNA expansion is altered in *top1* mutants. As any additional copy-number loss in critically short arrays is associated with grow defects or lethality (Takeuchi et al., 2003), the production of the lower ladder primarily reflects *de novo* events. Such events must occur at a substantial rate to be detectable by Southern blotting, given there is little further amplification by cell proliferation. Taken together, our data show that failure to form the tailed intermediate in absence of Top1 is associated with profound rDNA instability.

## DISCUSSION

Here, we describe a novel rDNA intermediate with a variably sized ssDNA-containing tail that originates from the promoter region of the 35S rRNA gene. Mutants lacking topoisomerase I fail to form detectable amounts of this intermediate and display a unique Fob1-independent form of rDNA instability.

Although the intermediate is detected at levels similar to replication structures in cycling wild-type cells, its origin and function in rDNA metabolism remain largely unclear. The intermediate is not the result of uncontrolled rDNA recombination, as its abundance does not increase in classic rDNA instability mutants, such as *sir2*. Instead, it likely reflects some form of normal rDNA metabolism. Given the high transcriptional output of the rRNA genes, it is possible that the intermediate arises as a function of the topological stresses of rRNA transcription. In line with this possibility, RNAPI accumulates across the 35S rRNA gene in *top1* mutant cells (El Hage et al., 2010). Thus, the intermediate could represent nicking events along the 35S gene as Top1 resolves stresses associated with the high level of rRNA transcription or the formation of RNA:DNA hybrids (El Hage et al., 2010). This model could explain why the transition from single-stranded to double-stranded DNA coincides with the 35S promoter. Arguing against this model is the fact that the reduced level of rRNA transcription in *rpa49, uaf30*, or *hmo1* mutants did not overtly affect the formation of the intermediate. Similarly, *trf4* and *hpr1* mutants, which accumulate RNA:DNA hybrids and show rDNA instability (El Hage et al., 2010, Luna et al., 2019, Merker and Klein, 2002), formed normal levels of the intermediate (**Supplemental Figure 2** and data not shown).

We also deem it unlikely that the intermediate reflects rolling-circle replication of ERCs. Although rolling-circle replication features long single-stranded tails (Garcillan-Barcia et al., 2022), the presence of the intermediate does not correlate with ERC levels: *fob1* mutants, which have strongly reduced ERC levels (Defossez et al., 1999), form wild-type levels of the intermediate, whereas *top1* mutants have increased ERC levels but no intermediate.

Intriguingly, fast migrating arcs of unknown function, similar to the one described here, are seen in 2D-gel analyses of maize streak virus DNA and *C. albicans* mitochondrial DNA (Gerhold et al., 2010, Erdmann et al., 2010). Maize streak virus replicates using rolling-circle replication as well as recombination-dependent mechanisms (Erdmann et al., 2010), whereas *C. albicans* mitochondrial DNA is linear and thought to exclusively use recombination-dependent replication (Gerhold et al., 2010). Thus, the tailed rDNA intermediate could potentially reflect recombination-dependent metabolism that occurs in wild-type cells. However, it is unclear how Top1 would interface with such recombination-dependent DNA synthesis. Although substantial transcription-independent Top1 cleavage activity occurs near the 35S promoter (Vogelauer and Camilloni, 1999, Krawczyk et al., 2014), homologous recombination is usually stimulated by DNA double strand breaks, rather than the nicks produced by Top1. The intermediate also does not noticeably change in mutants lacking known strand-invasion (*rad51 rad59*) or strand-annealing activities (*rad52 rad59*) (**Supplemental Figure 2**). Thus, the intermediate arises independently of the canonical homologous recombination machinery.

Regardless of its origin, our data suggest that failure to properly form the tailed intermediate in *top1* mutants is associated with substantial rDNA instability. This instability does not require Fob1 and thus contrasts with most known forms of rDNA instability, which are either partially or fully dependent on Fob1 (Weitao et al., 2003, Kaeberlein et al., 1999). Only very few mutants, including mutants defective in the Smc5/6 complex and mutants lacking the histone chaperone Asf1, are characterized by Fob1-independent rDNA instability (Moradi-Fard et al., 2021, Houseley and Tollervey, 2011). The instability of *smc5/6* mutants is likely linked to non-allelic homologous recombination resulting from the failure to move spontaneous DSBs outside of the nucleolus (Torres-Rosell et al., 2007), whereas the instability of *asf1* mutants requires DNA polymerases and thus involves DNA synthesis (Houseley and Tollervey, 2011). We found that *asf1* mutants exhibit normal formation of the tailed intermediate, but it remains possible that either Asf1 or Smc5/6 act downstream in guiding the proper processing or repair of this structure.

While investigating the instability of *top1* mutants we also uncovered an unexpected form of stepwise rDNA amplification that helps expand critically short rDNA arrays in a Fob1-independent manner. The fact that this laddering still occurs in *top1* mutants suggests that it does not require the controlled formation of the tailed intermediate. However, it is also possible that we failed to detect the tailed intermediate in *top1* mutants because the build-up of topological stress in *top1* mutants leads to distributed DNA damage, resulting in tailed intermediates forming stochastically across the rDNA instead of being focused at the 35S promoter. Formation of increased distributive DNA breakage would be consistent with the observation that strains with low rDNA copy number experience increased conflict between transcription and replication machineries (Takeuchi et al., 2003), which may be further exacerbated in *top1* mutants, given the higher density of polymerases observed across 35S genes in these mutants (El Hage et al., 2010).

Topoisomerase I is highly enriched in the nucleolus of many organisms and is thought to be important for supporting rRNA transcription (Muller et al., 1985, Huang et al., 2021, Fleischmann et al., 1984, Cha et al., 2012). However, topoisomerase I may also have additional, transcription-independent roles, as topoisomerase I enrichment in nucleoli precedes detectable enrichment of RNAPI in bovine embryos (Laurincik et al., 2000). Notably, amphibian oocytes, which display dramatic stage-specific rDNA amplification, exhibit strong nucleolar enrichment of a Top1 isoform (Gebauer et al., 1996). Thus, topoisomerase I may have functions in controlling rDNA metabolism and copy numbers outside of preventing transcription-associated DNA damage.

## MATERIALS AND METHODS

### Yeast strains and growth conditions

All strains used in this study were of the SK1 background, with the exception of the *cdc15-2* strain, which was congenic with SK1 (backcrossed 7x). The genotypes are listed in **Supplementary Table 1**. Unless otherwise noted, samples were collected from logarithmically growing cycling cultures. Cultures were seeded in YPD (YEP + 2% dextrose) at OD_600_ = 0.2 using saturated overnight cultures and grown for 4-5 hours at 30°C. To enrich cells in G1, alpha-factor peptide (GenScript) was added to a final concentration of 5 µg/ml for 1.5 hours. *cdc15-2* arrest/release was performed following a published protocol (Futcher, 1999). Briefly, an overnight culture was diluted to 600 ml YPD to an OD_600_ = 0.15 and grown at room temperature for 1 hour before shifting to 37°C in an air shaker to allow for slow accumulation of cells in telophase over 3.5 hours. The culture was then rapidly cooled to room temperature by temporarily placing the flask in an ice bath while shaking. The time when the flask was placed back on a shaker at room temperature was set as 0 min. Critically short arrays were produced as previously described (Kobayashi, 2006). Briefly, *fob1Δ* cells were transformed with a high-copy plasmid carrying a full rDNA copy with a mutation conferring resistance to hygromycin (Chernoff et al., 1994). Transformants were selected on hygromycin to identify clones that had lost most of their endogenous rDNA copies. We verified copy-number loss by pulsed-field gel electrophoresis and Southern blotting. Low-copy strains were mated and sporulated, and isolates that had lost the rDNA covering plasmid were immediately frozen for further manipulation. For sibling analysis, we marked a 13-repeat array with a *HIS3* marker (inserted next to *ARS1216*, <500bp to the left of the array) (Vader et al., 2011). This array was kept stable by deleting *FOB1* and by creating diploids in which the other array was of wild-type length. The diploids also carried a heterozygous deletion of *TOP1*. To initiate sibling analysis, diploids were sporulated, and tetrads scored to identify tetrads in which one sibling spore inherited the marked short array with wild-type *TOP1* and another sibling inherited the short array along with the *top1Δ* deletion. These sibling pairs were then analyzed for differences in rDNA size and ERC formation.

### Genomic DNA extraction

Genomic DNA was extracted as described previously (Mansisidor et al., 2018). Briefly, approximately 8.5 × 10^7^ cells were treated with 250 µg/ml zymolyase T100 (US Biological) in spheroplasting buffer (1 M sorbitol, 42 mM K_2_HPO_4_, 8 mM KH_2_PO_4_, 5 mM EDTA, 143 mM β-mercaptoethanol). Spheroplasts were lysed by addition of 0.5% SDS and proteins degraded using proteinase K (Roche). SDS was precipitated using 1M potassium acetate and removed by centrifugation. Nucleic acids were precipitated in ethanol, resuspended in TE (10 mM Tris pH 8.0, 1 mM EDTA), and incubated overnight with RNase A. DNA was purified using phenol/chloroform/isoamylalcohol (Roche), precipitated using isopropanol and resuspended in TE. Genomic DNA was digested for 2-3 hours at 37°C with StuI or SphI restriction enzymes (NEB), as indicated in the figure legends. For S1 treatment, samples were digested with StuI as above, ethanol-precipitated, then digested with S1 nuclease (Thermo Fisher Scientific) according to the manufacturer’s protocol.

### Gel electrophoresis

Standard gel electrophoresis of S1-treated samples was performed in a 0.7% agarose gel in 1X TBE buffer at 30V for 20 hours. Two-dimensional gel electrophoresis was performed in 0.4% SeaKem Gold agarose (Lonza) and 1X TBE in the first dimension. After equilibrating the gel in 1X TBE and 0.5 µg/ml ethidium bromide, lanes were cut, rotated 90°, embedded in 1% SeaKem LE agarose, and DNA separated in 1X TBE/0.5 µg/ml ethidium bromide in the second dimension.

### Pulsed-field gel electrophoresis

Approximately, 3 × 10^7^ cells per plug were embedded in 0.4% SeaPlaque low-melting-point agarose (Lonza) and in the presence of 75 mM EDTA. Cells were spheroplasted overnight at 37°C using 200 µg/ml zymolyase T100 in 500 mM EDTA and 10 mM Tris (pH 7.5). Cells were lysed by addition of 1% N-lauryl sarcosine and proteins digested with 0.4 mg/ml proteinase K overnight at 50°C. Proteinase K was inactivated by incubating plugs in CHEF TE (50 mM EDTA, 10 mM Tris, pH 7.5) with 1 mM phenylmethylsulfonyl fluoride at 4°C for 1 hour. Plugs were washed several times in CHEF TE to remove remaining detergent, equilibrated in CutSmart buffer and digested overnight with XhoI restriction enzyme (NEB), which does not cut in the rDNA. Plugs were embedded in 1% SeaKem LE agarose and rDNA fragments separated in 0.5X TBE in a BioRad CHEF-DR II instrument at settings: 5.1 V/cm, 23 h, 5-20 s switch-time ramp (for short rDNA fragments) or 6 V/cm, 60 s switch time for 15 h, followed by 90 s switch time for 9 hours (for large arrays).

### Southern blotting

We performed Southern blotting using alkaline transfer onto Hybond-XL (Cytiva) or ZetaProbe GT membranes (BioRad). Blots were labeled with ^32^P-labeled probes and signals detected using FujiFilm imaging plates and imaged using an Amersham Typhoon RGB instrument. Probes were made by nested PCR using the following internal primers: Probe A (18S; forward: 5’-CCAGAACGTCTAAGGGCATC-3’, reverse: 5’-TCGACCCTTTGGAAGAGATG-3’), Probe B (*NTS1*; forward: 5’-TCGCCGAGAAAAACTTCAAT-3’, reverse 5’-TGCAAAAGACAAATGGATGG-3’). All experiments were replicated at least twice in independent experiments either conducted on different days and/or using multiple yeast lines with identical genotypes (biological replicates; see Supplementary Table 1).

### Materials availability statement

Yeast strains created for this study are listed in Supplementary Table 1 and are available upon request.

## ACKNOWLEDGEMENTS

This work was supported by the National Institutes of Health [R01GM111715 and R01GM123035 to A.H.].

## AUTHOR CONTRIBUTIONS

Conceptualization, T.M., D.S., H.K. and A.H.; Investigation, T.M., D.S. and A.H.; Formal analysis, T.M., D.S., H.K. and A.H.; Writing – Original Draft, T.M. and A.H.; Writing – Review & Editing, T.M., D.S., H.K. and A.H.

## SUPPLEMENTAL MATERIAL

**Supplemental Figure 1.**
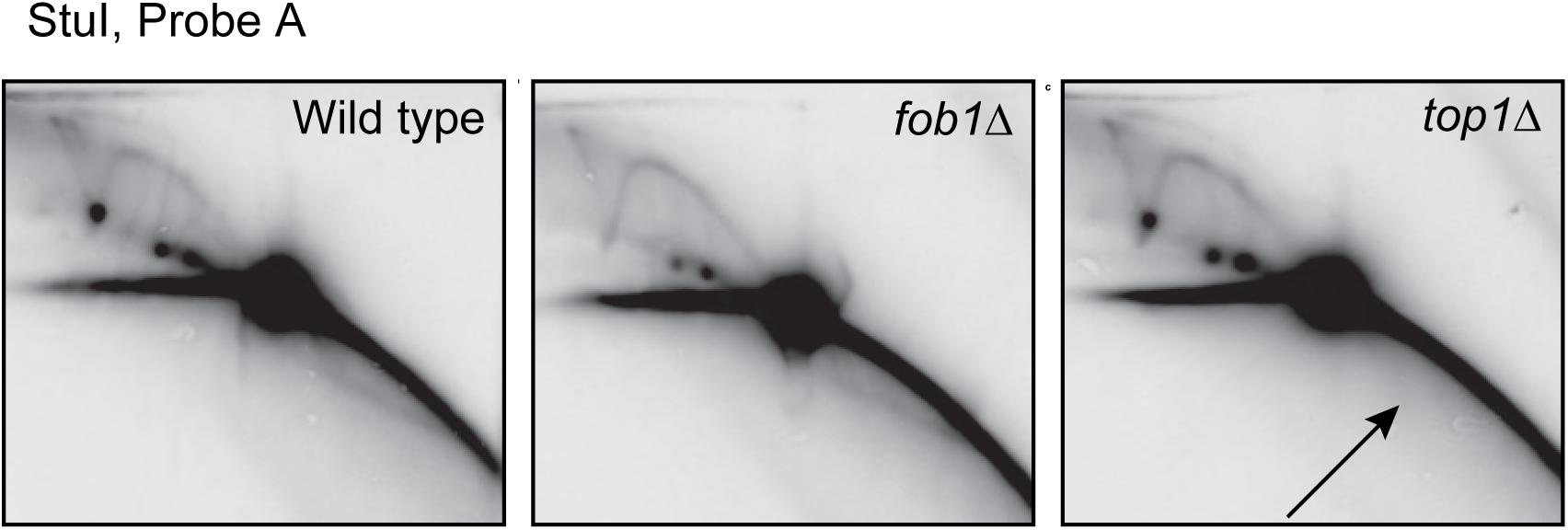
The tailed intermediate requires Top1 but is independent of Fob1. Two-dimensional gel electrophoresis and Southern analysis of a StuI rDNA fragment from wild type, *fob1Δ*, or *top1Δ* logarithmic cultures. Arrow indicates the absence of the tailed intermediate in *top1Δ* samples.

**Supplemental Figure 2.**
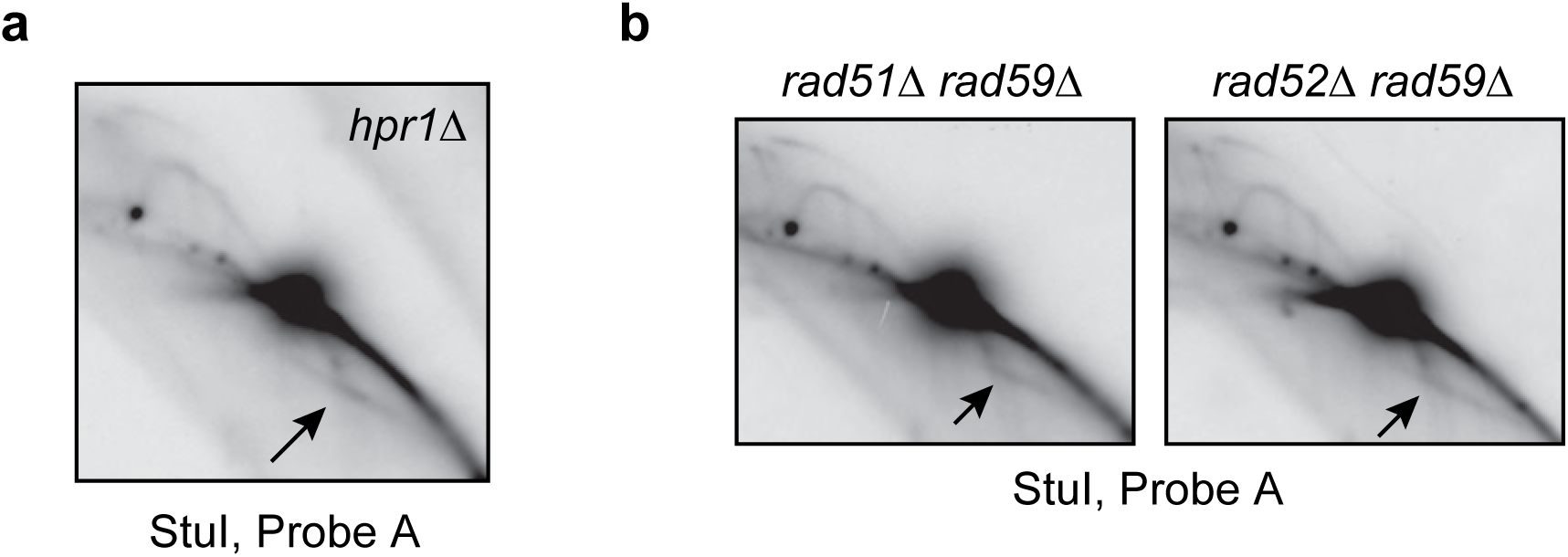
The tailed intermediate is unaffected in mutants with increased RNA:DNA hybrids or mutants defective for homologous recombination. Two-dimensional gel electrophoresis and Southern analysis of a StuI rDNA fragment from logarithmic cultures of (a) *hpr1Δ* mutants, which have elevated levels of RNA:DNA hybrids (Luna et al., 2019), and (b) *rad51Δ rad59Δ* double mutants or *rad52Δ rad59Δ* double mutants, which are defective in strand invasion and strand annealing, respectively. Arrows indicate the tailed intermediate.

**Supplemental Table 1.**
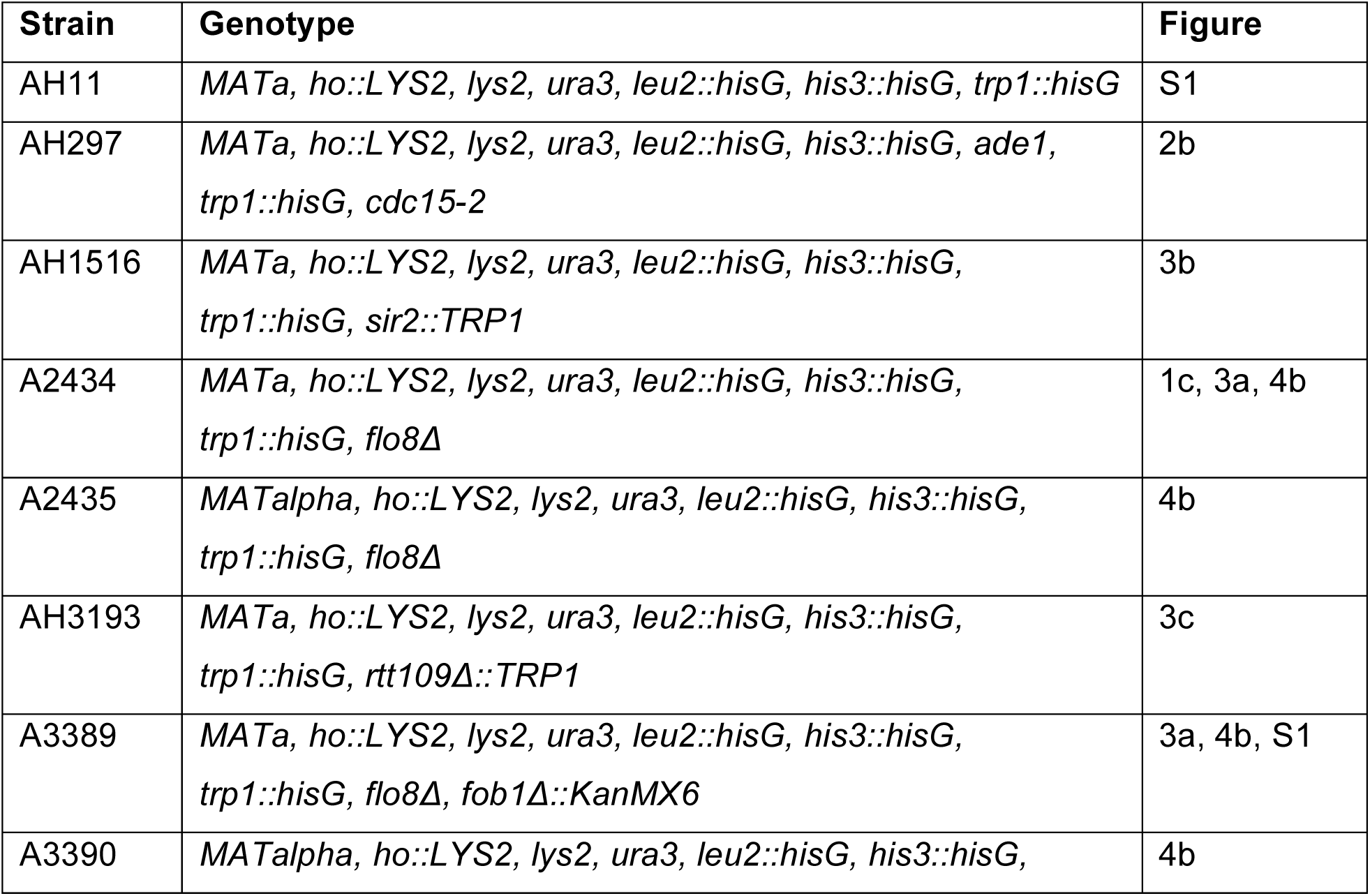
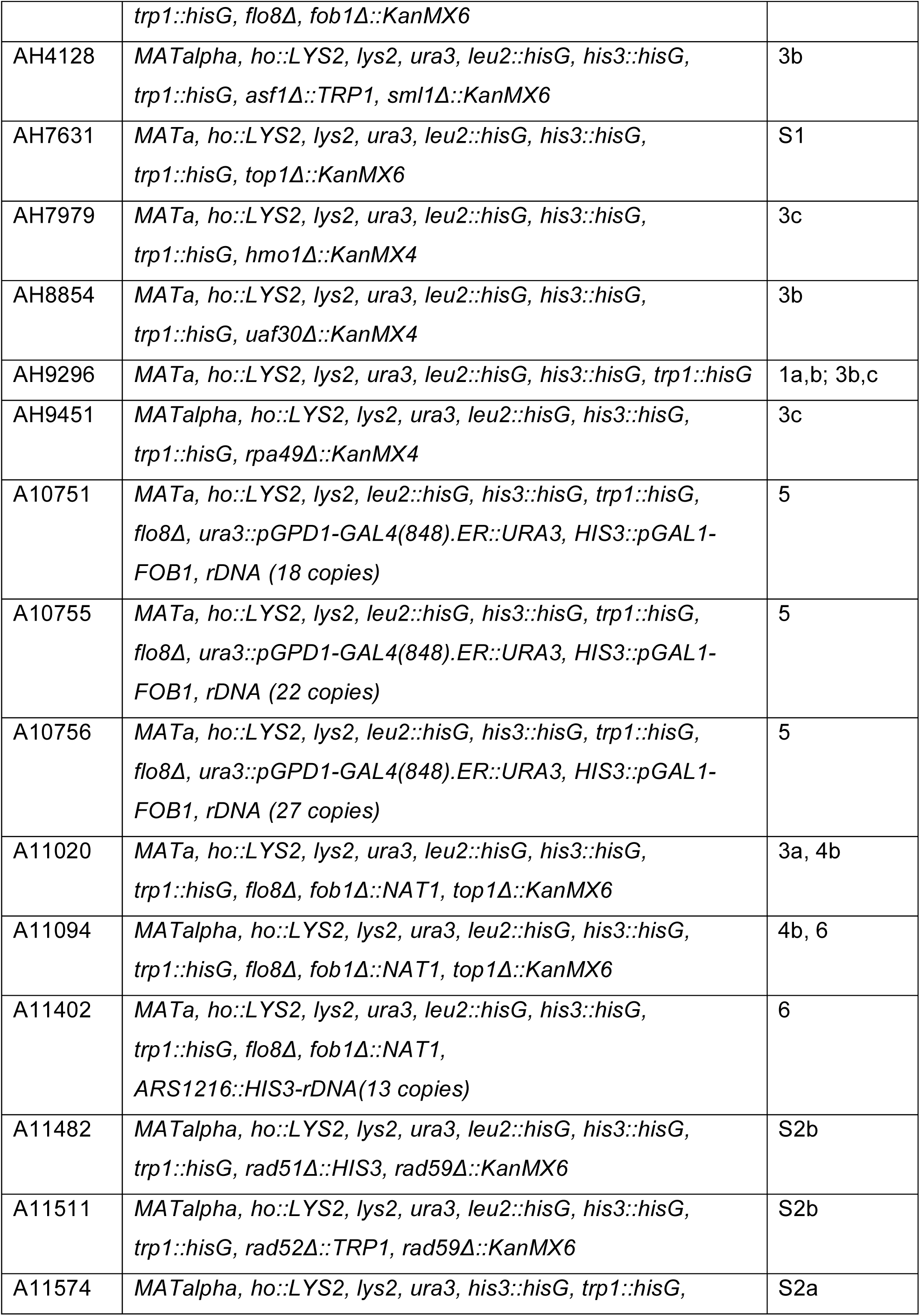
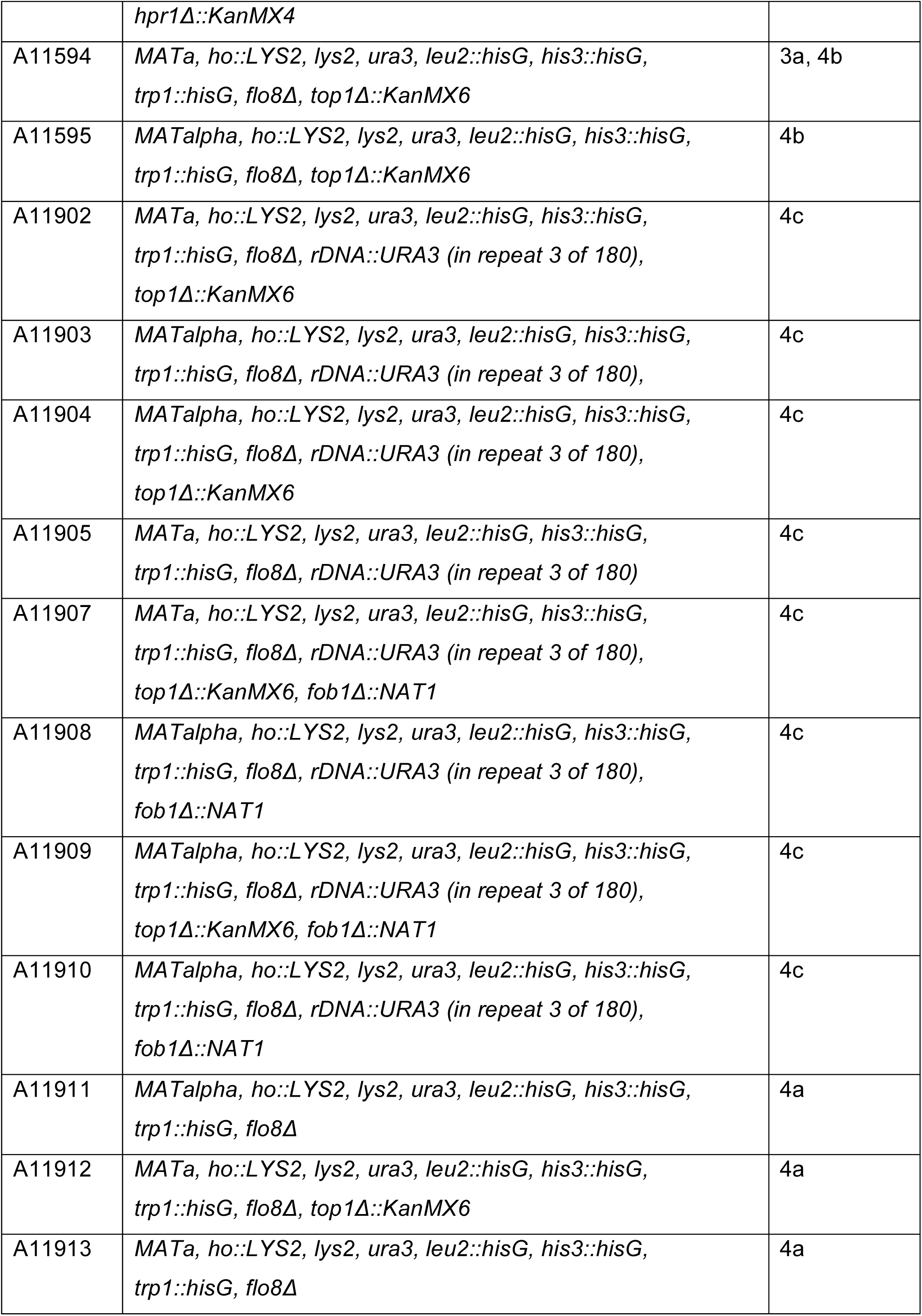
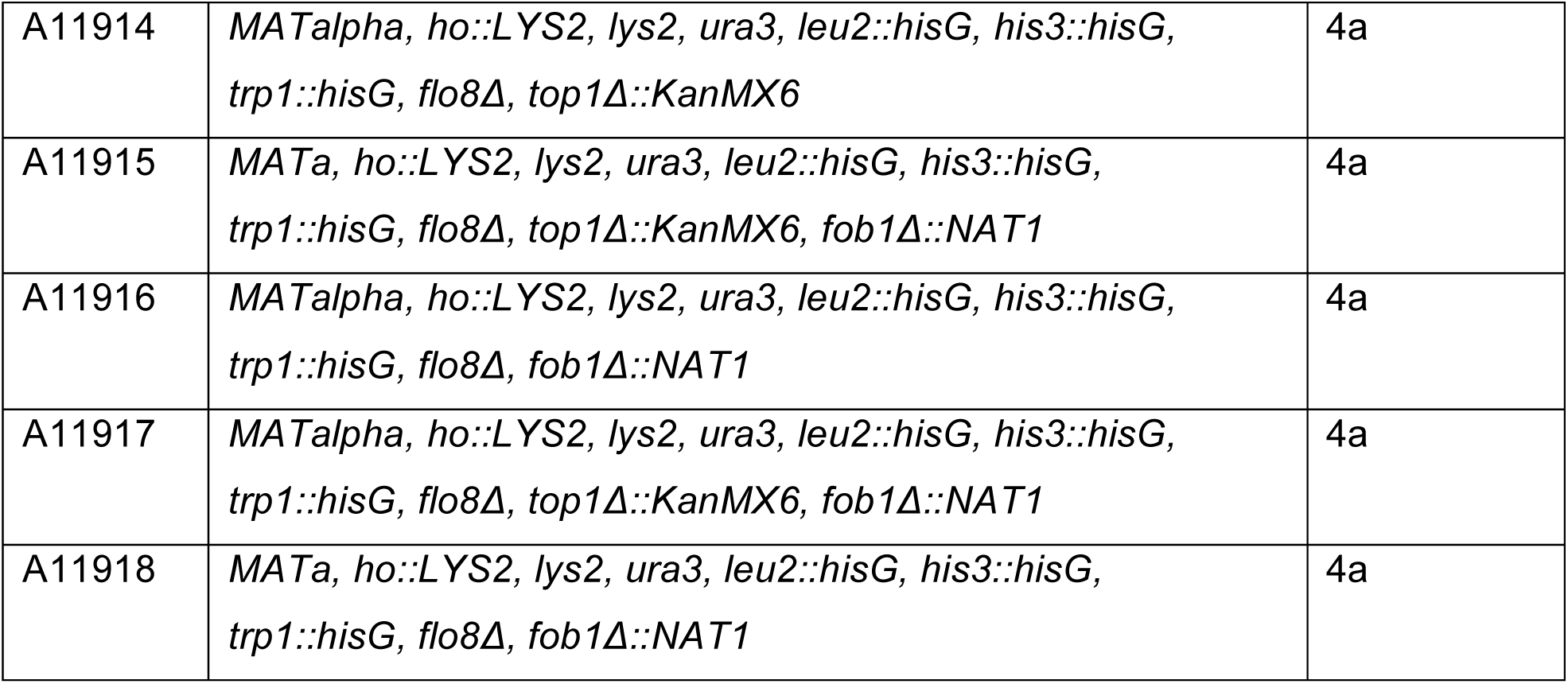
Strains used in this study.

